# Evolutionary dynamics of transposable elements following a shared polyploidization event in the tribe Andropogoneae

**DOI:** 10.1101/2020.03.05.978643

**Authors:** Dhanushya Ramachandran, Michael R. McKain, Elizabeth A. Kellogg, Jennifer S. Hawkins

## Abstract

Both polyploidization and transposable element (TE) activity are known to be major drivers of plant genome evolution. Here, we utilize the *Zea-Tripsacum* clade to investigate TE activity and accumulation after a recent shared polyploidization event. Comparisons of TE evolutionary dynamics in various *Zea* and *Tripsacum* species, in addition to two closely related diploid species, *Urelytrum digitatum* and *Sorghum bicolor*, revealed existing variation in repeat content among all taxa included in the study. The repeat composition of *Urelytrum* is more similar to that of *Zea* and *Tripsacum* compared to *Sorghum*, despite the similarity in genome size with the latter. Although the genomes of all species studied had abundant LTR-retrotransposons, we observed an expansion of the *copia* superfamily, specifically in *Z. mays* and *T. dactyloides*, species that have adapted to more temperate environments. Additional analyses of the genomic distribution of these *copia* elements provided evidence of biased insertions near genes involved in various biological processes including plant development, defense, and macromolecule biosynthesis. The lack of *copia* insertions near the orthologous genes in *S. bicolor* suggests that duplicate gene copies generated during polyploidization may offer novel neutral sites for TEs to insert, thereby providing an avenue for subfunctionalization via TE insertional mutagenesis.

## Introduction

Transposable element (TE) activation and accumulation generates significant genetic variation that can confer a range of effects on genome structure and function. As TEs carry ‘ready-to-use’ cis-elements, their insertions can impact gene regulation on a genome-wide scale by providing assorted regulatory elements to the adjacent genes. The new regulatory elements offered by inserted TEs can amplify and/or redistribute transcription factor binding sites, thereby creating new regulatory networks or even participate in re-wiring of pre-existing networks (Hènaff et al. 2014, Lavialle et al. 2013, Krupovic et al. 2014, Huang et al. 2016, Carmona et al. 2016, Joly-Lopez et al. 2016). Several empirical studies have demonstrated TE-induced phenotypic changes associated with domestication and/or diversification of cultivated plants, including rice, maize, wheat, soybean, melon, and palm (Naito et al 2009, Fernandez et al 2010, Studer et al. 2011, Uchiyama et al. 2013, Sanseverino et al. 2015, Ong-Abdullah et al. 2015, Lu et al. 2017). Indeed, TE-related polymorphisms are largely responsible for phenotypic variation in many agronomically important crops, demonstrating their importance in creating the genetic variability that contributes to plant genome evolution.

Hybridization, polyploidy, and stress are considered the primary triggers of transposable element movement (Steward et al. 2000, Kalendar et al. 2000, Madlung et al. 2005, Ungerer et al 2006, Ito et al. 2011, Cavrak et al 2014, Bardil et al. 2015, Guo et al. 2017). Flowering plants are known to tolerate hybridization and polyploidy, both of which have promoted species diversification (Payseur and Rieseberg, 2016, Soltis et al. 2016, Goulet et al. 2017). These phenomena result in TE mobilization leading to local mutations and genome size changes (Liu and Wendel 2000; Josefsson et al. 2006; Ungerer et al. 2006; Kawakami et al. 2010; Parisod et al. 2010; Piednoël et al. 2013). Furthermore, such bursts of TE activity result in insertional polymorphisms, often with deleterious effects on genome function; however, these effects could be nullified or shielded via gene duplication in polyploid genomes. Although the precise mechanism(s) that induce TE mobility in hybrids and polyploids is unclear, it is speculated that such TE reactivation in response to genomic stresses could be due to incompatible suppression machinery between the two donor genomes, or that unknown mechanisms are in place that reduce genomic methylation under general stress conditions (Ha et al. 2009, Yaakov and Kashkush 2012, An et al. 2014, Senerchia et al. 2014, DeFraia and Slotkin 2014, Ågren et al. 2016).

Previous studies of polyploidy in *Zea* have revealed evidence for a whole genome duplication (WGD) event at or shortly after the origin of grasses (Paterson et al. 2004), followed by another, more recent, WGD in the *Zea* history that promoted the origin of the *Zea-Tripsacum* clade (Estep et al. 2014; McKain et al. 2018). Diversifying from a common ancestral allotetraploid (n=20), both the *Zea* (n=10) and *Tripsacum* (n=18) genomes differentially responded to the diploidization process (Swignova et al. 2004, Schnable et al. 2009, Schnable & Freeling, 2011). In addition to these chromosomal rearrangements, there is also evidence for retrotransposon invasion post divergence in both *Zea* and *Tripsacum* (Gaut et al. 2000). Hence, being divergent descendants of a common allopolyploid ancestor, the *Zea-Tripsacum* clade is a good model system to understand various evolutionary processes including the contribution of TEs to polyploidy, rediploidization, and species diversification.

Here, we describe TE activity and contribution to genome diversity in the *Zea-Tripsacum* clade that has undergone a recent shared polyploidization event (**Figure 1A**). We included two diploid relatives, *Urelytrum digitatum* and *Sorghum bicolor*, which provide an opportunity to explore TE-associated evolutionary events induced by hybridization and genome doubling. By using clustering analysis, we have characterized the repetitive landscape in three *Zea* and three *Tripsacum* species (post allopolyploidization) and compared these results with that of the diploid relatives (pre-allopolyploidization). Our findings suggest recent and post-divergence activity of TEs in *Zea* and *Tripsacum* with lineage-specific expansion of *copia* elements in *Z. mays* and *T. dactyloides*. Insertional biases of these newly expanded *copia* elements near developmental and defense genes may have influenced the evolution of the *Z. mays* and *T. dactyloides* genomes during adaptation to temperate climates.

**Figure 1.**
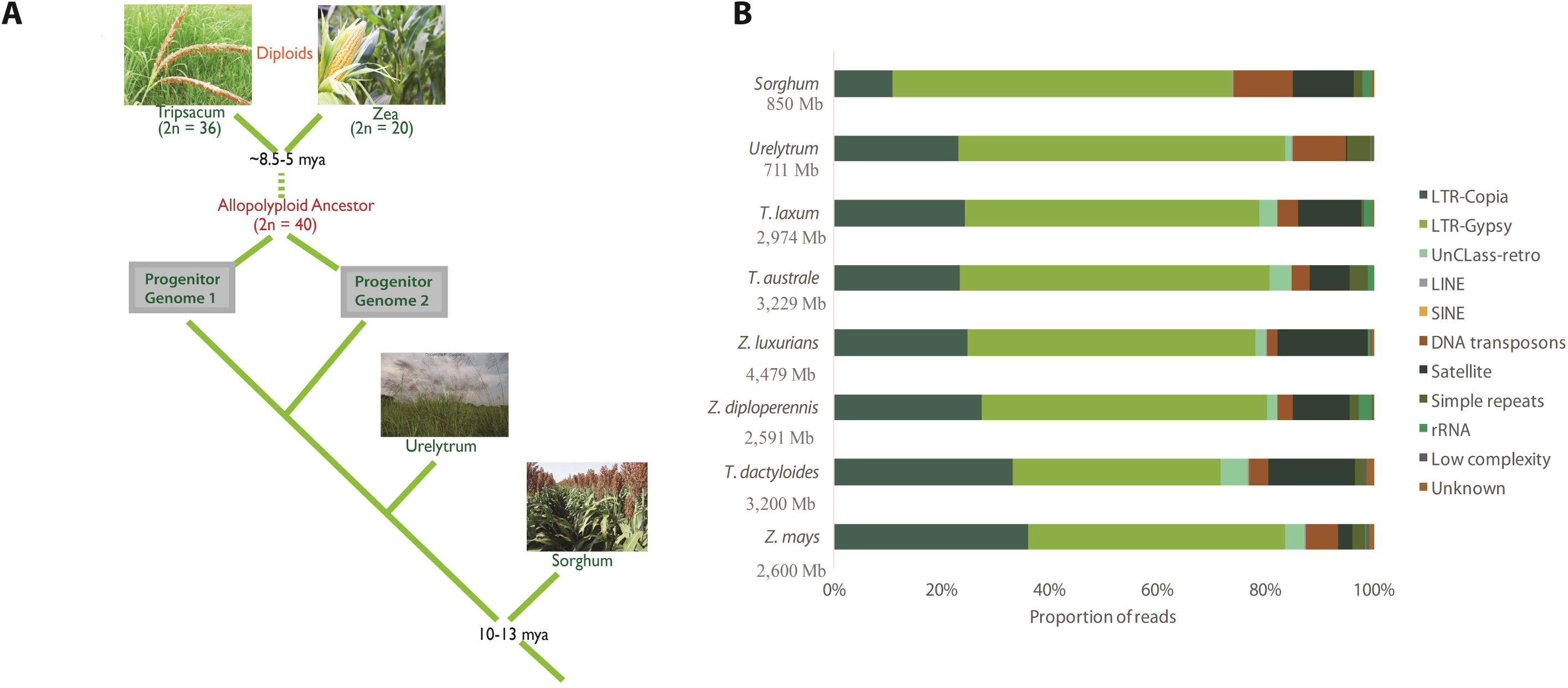
**A.** The evolutionary relationships of selected grass species, indicating polyploidization and species divergence. **B**. Proportional repeat composition. Genome size in Mb shown for each species in the y-axis. An expansion of *copia* families is observed in both cultivated *Z. mays* and *T. dactyloides* compared to related sister species. *Sorghum* displays a predominance of *gypsy* elements with a low level accumulation of *copia* families compared to *Zea-Tripsacum-Urelytrum* genomes.

## Materials and methods

### Plant material sources and Illumina sequencing of DNA

The following eight panicoid grasses were used in this study: *Zea mays L., Z. diploperennis Iltis, Doebley & R.Guzmán, Z. luxurians (Durieu & Asch.) R.M.Bird, Tripsacum dactyloides (L.) L., T. laxum Nash, T. australe Cutler & E.S.Anderson, Urelytrum digitatum K.Schum., and Sorghum bicolor (L.) Moench*. Short-read sequence data for *Zea mays* (SRS291653), *Zea luxurians* (SRR088692), *Tripsacum dactyloides* (SRS302460), and *Sorghum bicolor* (SRR5271056) were downloaded from the NCBI short read archive (Chia et al. 2012, Tenaillion et al. 2011, McCormick et al. 2018). Genomic short-read sequences of *Zea diploperennis* (XXXXXX – awaiting for an ID number from the short read archive), *Tripsacum laxum* (MIA34792), *Tripsacum australe* (MIA34499) and *Urelytrum digitatum* (SM3109) were obtained from Dr. Elizabeth Kellogg, Donald Danforth Plant Science Center, St. Louis, Missouri. See Supplementary Table 1 for more information on genome sequencing.

### Identification of TE families

Sequences were quality trimmed with Trimmomatic v0.33 (Bolger et al. 2014) using a sliding window of 4:25 and minimum length of 50 bp. Graph-based clustering of quality-trimmed reads was performed with RepeatExplorer, a pipeline designed to identify repeats from NGS reads (Novak et al. 2013). RepeatExplorer employs a clustering algorithm that quantifies similarities between all sequence reads and produces a graph that consists of nodes (sequence reads) and edges (connecting overlapping reads). Nodes are frequently connected to one another if they pass a threshold of 90% similarity over at least 55% of the sequence length, representing individual repetitive families.

Species-specific clustering analysis provides information regarding repeat quantities by reporting the number of reads per cluster, which can then be used to estimate the genome space occupied by each particular repeat, i.e., (total length of each cluster (in Mb) x genome size (in Mb)) / total length of all clusters (in Mb) (Kelly et al. 2015, Ramachandran et al. 2016). For species-specific clustering, three million reads (approximately 0.2x to 0.5x genome coverage) were sub-sampled from each dataset and processed to the format required by RepeatExplorer. Subsequently, all of the processed reads from all species were concatenated into one combined dataset, and the RepeatExplorer clustering was repeated in order to facilitate comparative analysis. All clusters were annotated using the Viridiplantae RepeatMasker library and categorized into repeat families. A plot representing interactions between repeat clusters among species was created using UpSetR (Lex et al. 2014).

### Quantitative analysis of TE activity using molecular clock analysis

To estimate the timing of TE activity in each lineage, species-specific LTR sequences were extracted from each LTR-retroelement cluster. These species-specific reads were assembled using the Geneious *de novo* assembler to obtain a consensus sequence (Kearse et al. 2012). A grass-specific database was then used to extract LTRs from each consensus contig (blastn, e-value 1e-10, 85% identity). The best match for each species was chosen and the corresponding hit region was extracted using BEDTools v2.17.0 (Quinlan and Hall 2010).

To calculate LTR divergence (a rough measurement to estimate the age of a specific retrotransposon family), reads that were used for *de novo* assembly were mapped to the consensus LTR sequence using the Geneious reference genome assembler. The percent identity of each read mapped to its respective LTR consensus sequence was derived from the reference alignment. Using a grass specific transposable element substitution rate of 1.3 × 10^−8^ per site per year (Ma and Bennetzen, 2004), we estimated the activity of each major TE family in each species.

### Genomic distribution of *copia* retroelements

To test whether the elements that have expanded in select species demonstrate an insertional bias, Illumina paired-end reads from *Z. mays* were mapped to a library consisting of *Z. mays copia* clusters assembled by RepeatExplorer and to a filtered gene set containing the protein-coding genes from the *Z. mays* reference genome. Reference mapping of paired-end reads to the library was carried out using BWA aln v0.7.12 (Li and Durbin, 2009) with the following optimized parameters: using the first 12 bases as a seed (-l 12), a maximum edit distance of four (-n 4), a maximum edit distance of two for the seed (-k 2), up to three gap openings (-o 3), a maximum of three gap extensions (-e 3), a mismatch penalty of two (-M 2), a gap opening penalty of six (-O 6), and a gap extension penalty of 3 (-E 3) (Mascagni et al, 2015). The results were used to generate a SAM file via the BWA “sampe” module, and then converted to a BAM file using SAMtools v1.9 (Li et al, 2009). A *copia* element was considered near a gene if one of the paired-end read mapped to a *copia* element and the other to a gene. Genes near *copia* elements were further analyzed for their presence in gene-dense or gene-poor regions by determining the number of TEs present within various distances (1 kb, 5 kb, and 10 kb) both upstream and downstream of genes using BEDTools v2.17.0 (Quinlan and Hall 2010). A similar analysis of *copia* insertions was performed for the *S. bicolor* (BTx623) genome using a library consisting of *S. bicolor copia* clusters and a filtered gene set containing *S. bicolor* protein coding genes.

As *Tripsacum* lacks a reference genome assembly, we performed *copia* analysis for the *T. dactyloides* genome using *Z. mays* reference protein coding genes. *T. dactyloides* paired-end reads that matched to *copia* clusters were identified using BLASTn v2.2.28+ (e-value = 1e-10, percent identity = 80). Identified reads were extracted and mapped to a library consisting of *copia* contigs and *Z. mays* protein coding genes as described above. Paired-end reads that are discordantly mapped to a repeat and a gene were extracted from BAM files, and genes were annotated via the available reference gene annotation file.

### Phylogenetic analysis of retroelement families

To assess the evolutionary relationships of the shared *gypsy* and *copia* families, the reverse transcriptase (RT) and integrase (INT) amino acid domains were used for phylogenetic analysis. RepeatExplorer clusters were filtered for LTR-*gypsy* and *copia* elements with RT and INT domain blastx hits. RT reads were extracted from each cluster using the blastx output file and placed in separate genome-specific files. The reads were assembled for each cluster using the Geneious *de novo* assembler (Kearse et al. 2012). The resulting contigs were then confirmed to contain reverse transcriptase domains using blastx against the Cores-RT database (Llorens et al. 2011). RT sequences were then combined into a final query file for further analysis. The same analysis was performed for INT reads using Cores-INT database (Llorens et al. 2011).

Rpstblastn (e-value = 1e-10) was performed for the sequence dataset against the Conserved Domain Database (Marchler-Bauer et al. 2015) to identify and extract conserved regions. The best hits for each sequence were extracted, and the filtered blast output was converted to three-column bed format with matching coordinates for each hit. BEDTools v2.170 (Quinlan and Hall 2010) was used to extract the conserved regions (∼540 bp for the RT domain and ∼340 bp for the INT domain). The correct open reading frame from each sequence was identified using ORFfinder. All amino acid sequences were globally aligned with MUSCLE v3.8.31 (Edgar 2004). Alignments were manually inspected and adjusted in Bioedit v7.3.5 (Hall, 1999). The optimal model of amino acid substitution for each alignment was estimated using Prot-test v3.4.5 according to the Akaike Information Criterion (AIC) (Abascal et al. 2005). In all cases except RT-*copia*, the best model selected was LG+G (Le and Gascuel. 2008). Blosum62+G was chosen as the optimal model for RT-*copia* (Henikoff and Henikoff. 1992). Likelihood analyses with 1,000 bootstrap replicates were performed in RAxML v.8.2.9 (Stamatakis et al. 2008) using the best model for each alignment. Bayesian analysis of alignments was performed in MrBayes v3.2.6 using rates=gamma and the respective substitution model (Ronquist and Huesenbeck. 2003). Two independent MCMC runs of 10 million generations were performed, sampling each run every 1,000 generations until convergence, with the potential scale reduction factor close to 1. All trees were visualized using FigTree v1.4.0.

## Results

### Single-species clustering analysis

To evaluate repeat content with respect to genome size, we performed a separate clustering analysis for each species. Individual clustering allows the maximum number of reads to assemble, which increases the accuracy of the repeat estimates. We estimated the quantities of each repeat family in the genome using the following equation: (total length of each cluster (in Mb) x genome size (in Mb)) / total length of all clusters (in Mb) (Macas et al. 2015). The estimated repeat compositions are shown in **Table 1**.

**Table 1:**
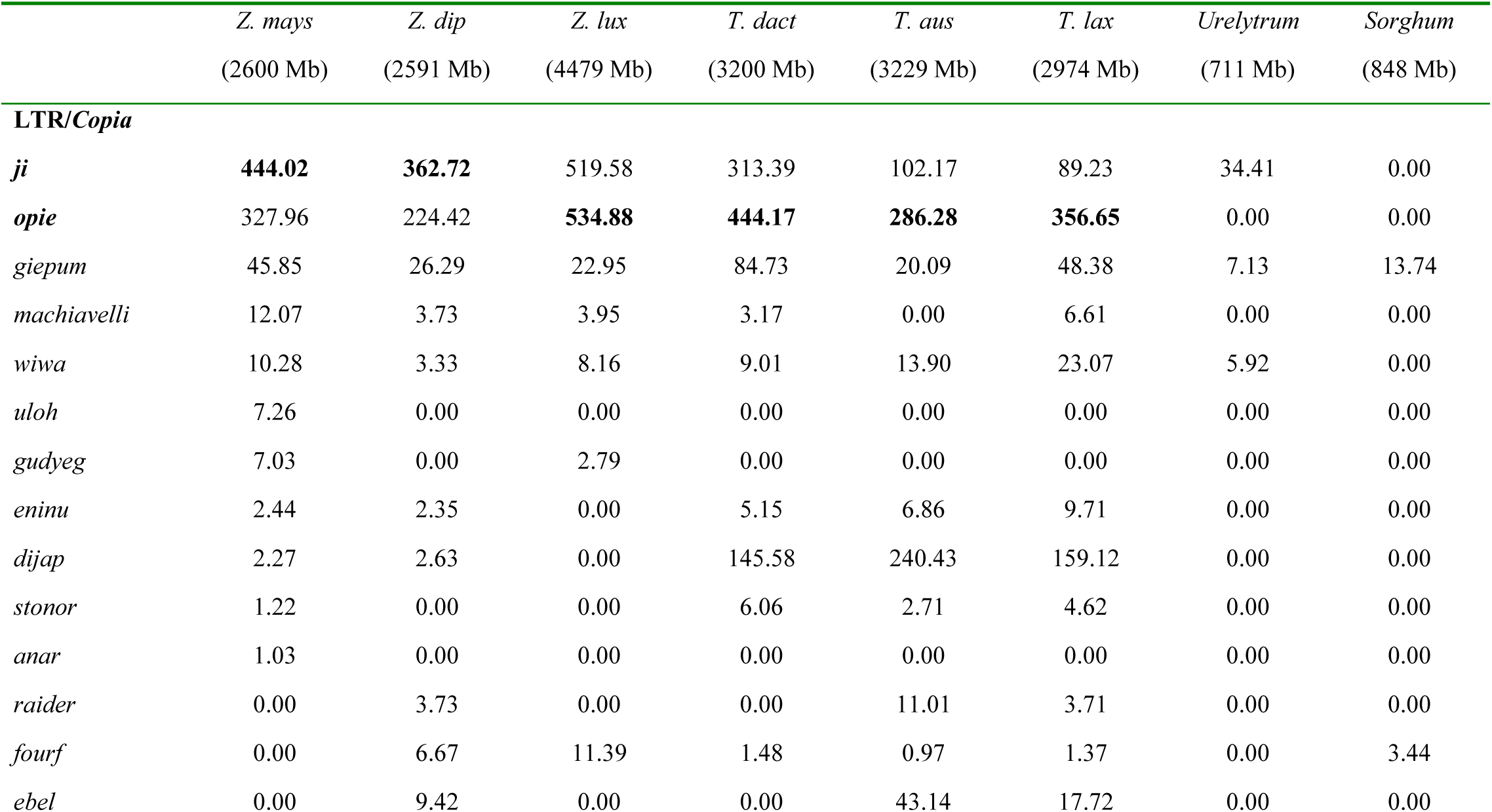
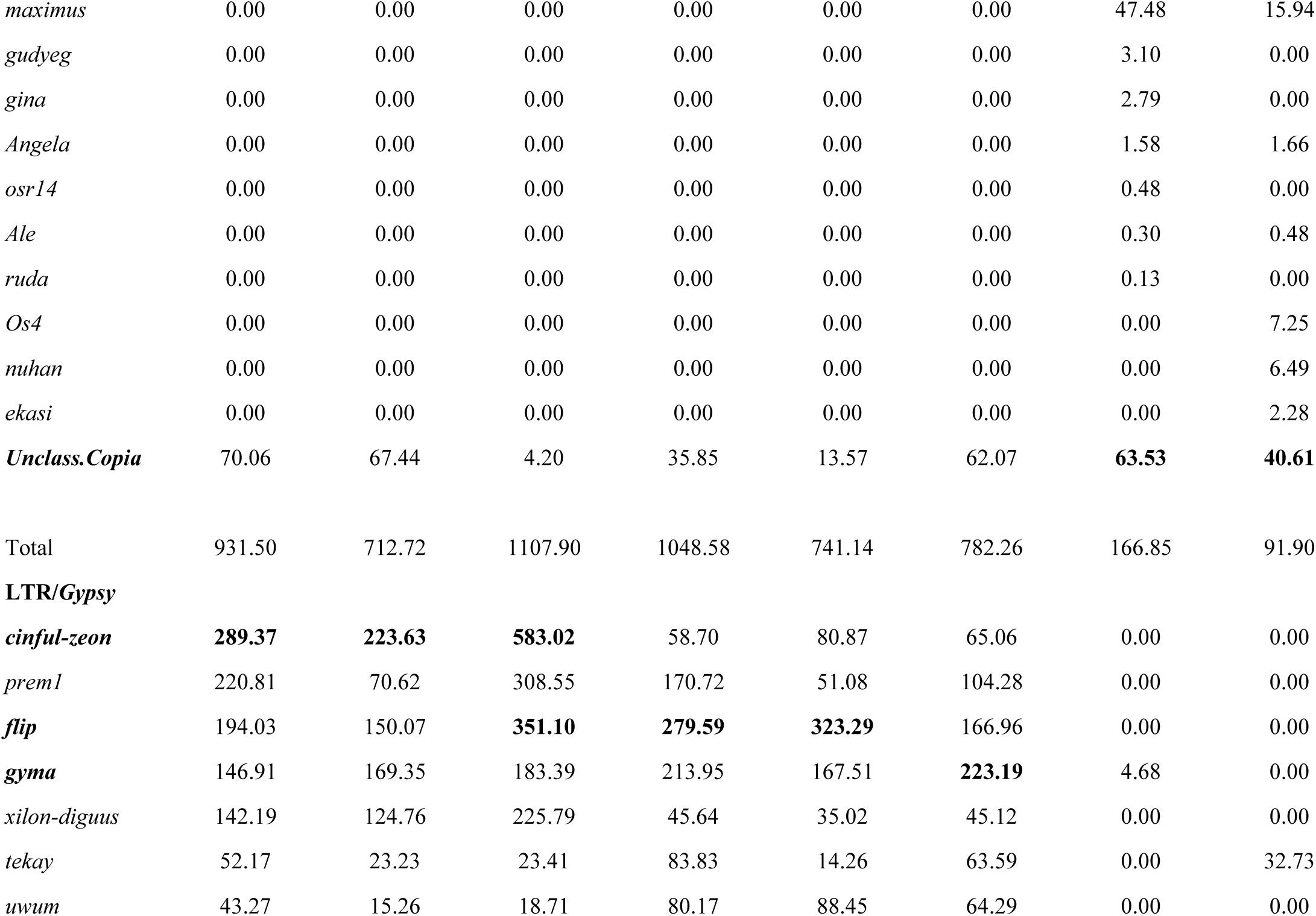
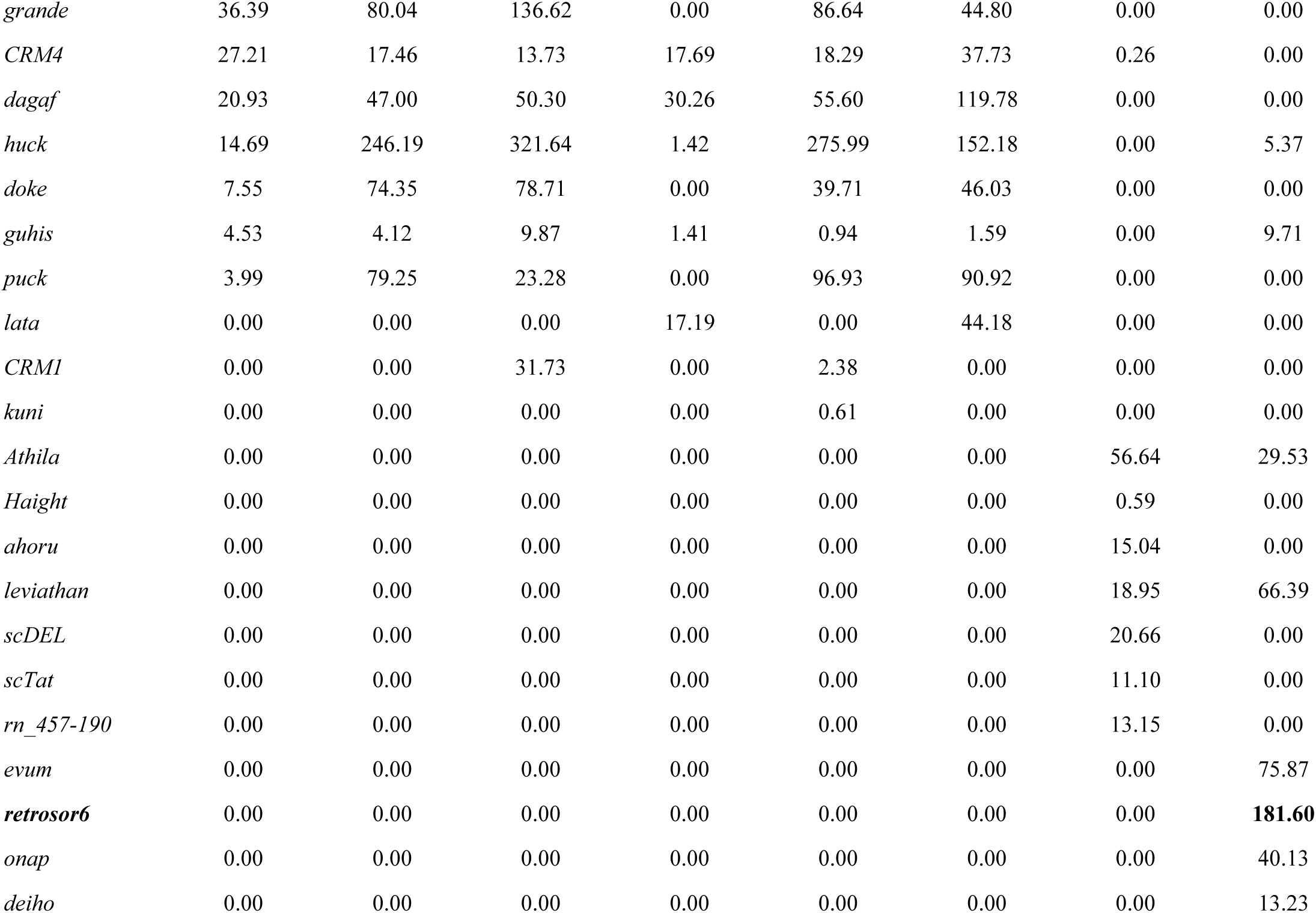
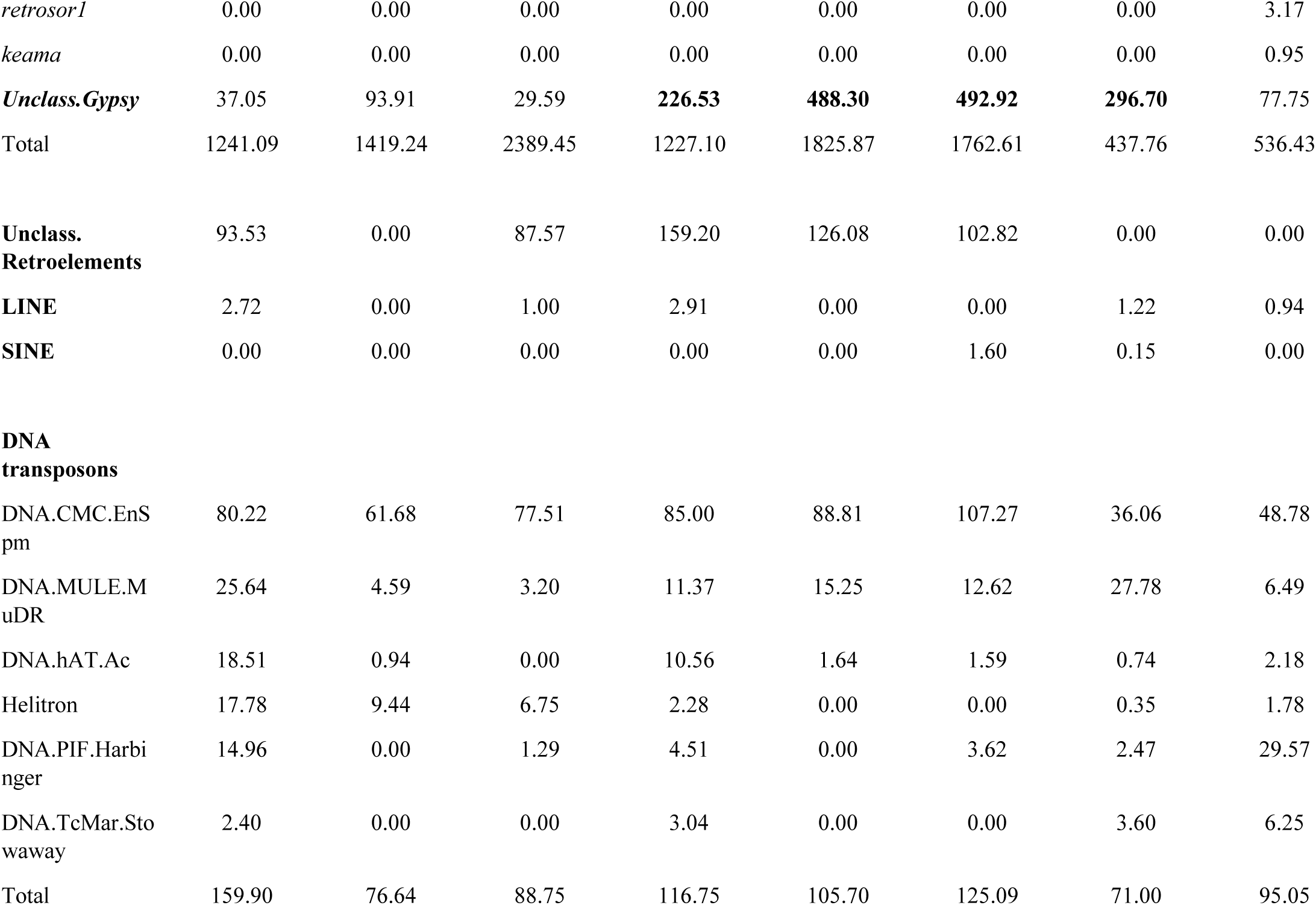
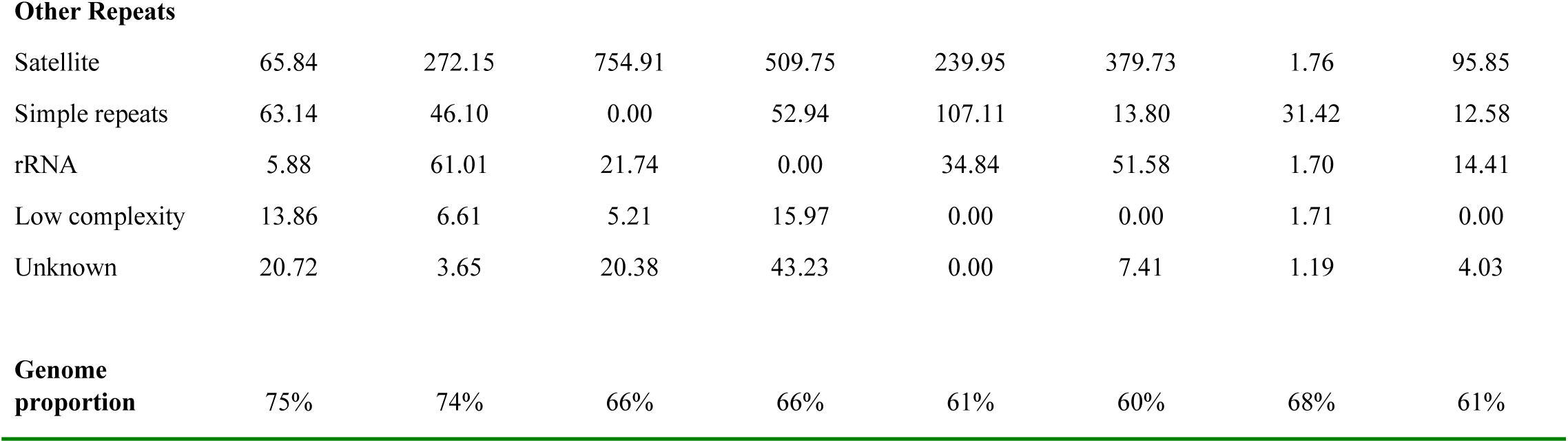
Global repeat composition (in Mb) of species with respect to genome size. Genome size (in Mb) for each species is given below each species name. Estimated repeat content (in Mb) for each repeat family is listed below using individual repeat clustering analysis. Bold text represents the most abundant families in each genome.

As expected, LTR-retrotransposons are the most abundant repeat in all eight species. Although all *Zea* species used in this study are diploid and contain the same number of chromosomes, the genome size of *Z. luxurians* (∼4,479 Mb) is nearly double the size of the other two *Zea* species (∼2,600 Mb). From the clustering analysis, *copia* elements were found to contribute approximately 710 Mb, 930 Mb, and 1,110 Mb to the *Z. diploperennis, Z. mays* and *Z. luxurians* genomes, respectively. *Gypsy* elements account for ∼1,240 Mb and 1,420 Mb of the *Z. mays* and *Z. diploperennis* genomes, respectively, whereas ∼2,390 Mb of *Z. luxurians* genome is comprised of *gypsy* elements (**Table 1**). The greater repeat abundance in *Z. luxurians* correlates with its larger genome size. The *Tripsacum* species have genomes of similar size (∼3,100 Mb) and chromosome number (2n=36), in which *T. laxum* possesses the smallest genome (2,974 Mb). *Copia* elements occupied ∼740 and 780 Mb in *T. australe* and *T. laxum* genomes, in contrast to 1,050 Mb in the *T. dactyloides* genome. Approximately 1,760 Mb and 1,825 Mb is composed of *gypsy* elements in *T. laxum* and *T. australe*, respectively, whereas the *T. dactyloides* genome contains 1,230 Mb of *gypsy* elements. *Gypsy* elements contributed to more of the genome space (∼53-59%) compared to *copia* (22-28%) in all *Zea* and *Tripsacum* species except for *Z. mays* and *T. dactyloides*, where both *gypsy* and *copia* were nearly equally abundant (**Figure 1B**). *Urelytrum* and *Sorghum* contain ∼167 Mb and 92 Mb of *copia*, and 438 Mb and 536 Mb of *gypsy* elements, respectively.

DNA transposons were found to contribute only 2-6% to the *Zea* and 2-3% to the *Tripsacum* genomes, in contrast to 10-11% in *Urelytrum* and *Sorghum* (Figure 1B). Other groups of repeat elements such as satellite repeats made up a significant fraction of the genome in several species. Approximately 755 Mb of the *Z. luxurians* and 510 Mb of the *T. dactyloides* genomes were occupied by satellite repeats. Although *Urelytrum* and *Sorghum* contain genomes of similar size, the former is composed of only 1.76 Mb of satellite DNA whereas the latter contained ∼100 Mb of satellite DNA.

### Repeat families and their contribution to genome size

From the individual repeat clustering analysis, we identified 24 *copia* and 30 *gypsy* families. Among the 24 *copia* families, *Ji* was the most abundant family in both *Z. mays* (444 Mb) and *Z. diploperennis* (363 Mb), whereas *Opie* was the most abundant in *Z. luxurians* (535 Mb) and in all of the *Tripsacum* species (**Table 1**). Disproportionately large increases are observed for *Ji* and *Opie* in *Z. mays* and *T. dactyloides*, in agreement with the overall increase of the *copia* superfamily observed for these two species. Interestingly, *Dijap* was estimated at 146-240 Mb in *Tripsacum*, but contributed very little to the genome size of *Zea* (0-2 Mb), indicating lineage-specific accumulation of this family in the *Tripsacum* genus.

Among the *gypsy* families, *Cinful-Zeon, Prem1, Flip, Gyma, Huck and Xilon-Diguus* were abundant in both *Zea* and *Tripsacum*. The *Cinful-Zeon* family ranges from 224 - 583 Mb in *Zea* and is positively correlated with genome size, with the greatest abundance in *Z. luxurians*; however, this family contributes only ∼70 Mb to the genomes of the *Tripsacum* species. This is also true for the *Xilon-Diguus* family, with estimates ranging from 125 - 226 Mb in *Zea* and ∼42 Mb in *Tripsacum*, indicating amplification in the lineage leading to *Zea* after divergence from *Tripsacum*. The *Huck* family is estimated at 246 Mb in *Z. diploperennis*, 321 Mb in *Z. luxurians*, 152 Mb in *T. laxum*, and 276 Mb in *T. australe*; however, *Huck* occupies only ∼15 Mb of the *Z. mays* genome and ∼1.4 Mb of the *T. dactyloides* genome. Similarly, elements such as *Doke* and *Puck* were noticeably more abundant in *Z. diploperennis, Z. luxurians, T. laxum*, and *T. australe*, suggesting independent lineage-specific loss of these families in these *Z. mays* and *T. dactyloides*.

There were 13 *gypsy* families that were specific to *Urelytrum* and/or *Sorghum. Athila* and *Leviathan* were relatively abundant (∼19 – 66 Mb) and were identified in both *Urelytrum* and *Sorghum*. Apart from these two families, the remaining 11 *gypsy* families were predominantly present in *Sorghum*, but present in low copy number in *Urelytrum*, or absent altogether. For example, *Retrosor6* is estimated at ∼180 Mb in the genome of *Sorghum* but is completely absent in all other species; however, there are a large number of unclassified *gypsy* elements in *Urelytrum* (**See Table 1**). Although we used a grass specific database to annotate the elements, the majority of this repeat content could not be annotated, suggesting the presence of species-specific repeats and retroelements.

### Comparative clustering analysis

We performed comparative repeat analysis by simultaneously clustering reads from all eight species in order to identify repeat families that are shared between multiple species and to determine their fate during Andropogoneae evolution, especially during the divergence of *Zea* and *Tripsacum*. This analysis resulted in four major cluster configurations, for which examples are shown **in Figure 2A-D. Figure 2A** shows an example of a cluster (2: *Prem1*, LTR-*gypsy*) in which the repeat family is common to all species. In this example, most of the reads from *Zea* and *Tripsacum* are tightly clustered, and reads from *Sorghum* and *Urelytrum* are peripherally connected, as would be expected based on their evolutionary relationships. Cluster 6 (*Opie*, LTR*-copia*) is an example of genus-specific repeat accumulation, where sequences are shared between *Zea* and *Tripsacum* but absent in *Urelytrum* and *Sorghum* (**Figure 2B**). In Cluster 21 (*Flip*, LTR-*gypsy*), the graph indicates three separate groups: one composed primarily of *Z. mays* and *T. dactyloides* [top], one composed primarily of *Z. diploperennis* and *Z. luxurians* [right], and one composed primarily of *T. australe* and *T. laxum* [left]). This type of configuration indicates post-divergence species-specific accumulation independently in *Z. mays* and *T. dactyloides* with minimal transpositional activity in their sister species (**Figure 2C**). Finally, cluster 64 (*Angela*, LTR-*copia*) is an example of a tightly knitted graph in a circular arrangement shared among all eight species, demonstrating the conserved nature of ancient *Angela* elements across all included taxa (**Figure 2D**).

**Figure 2.**
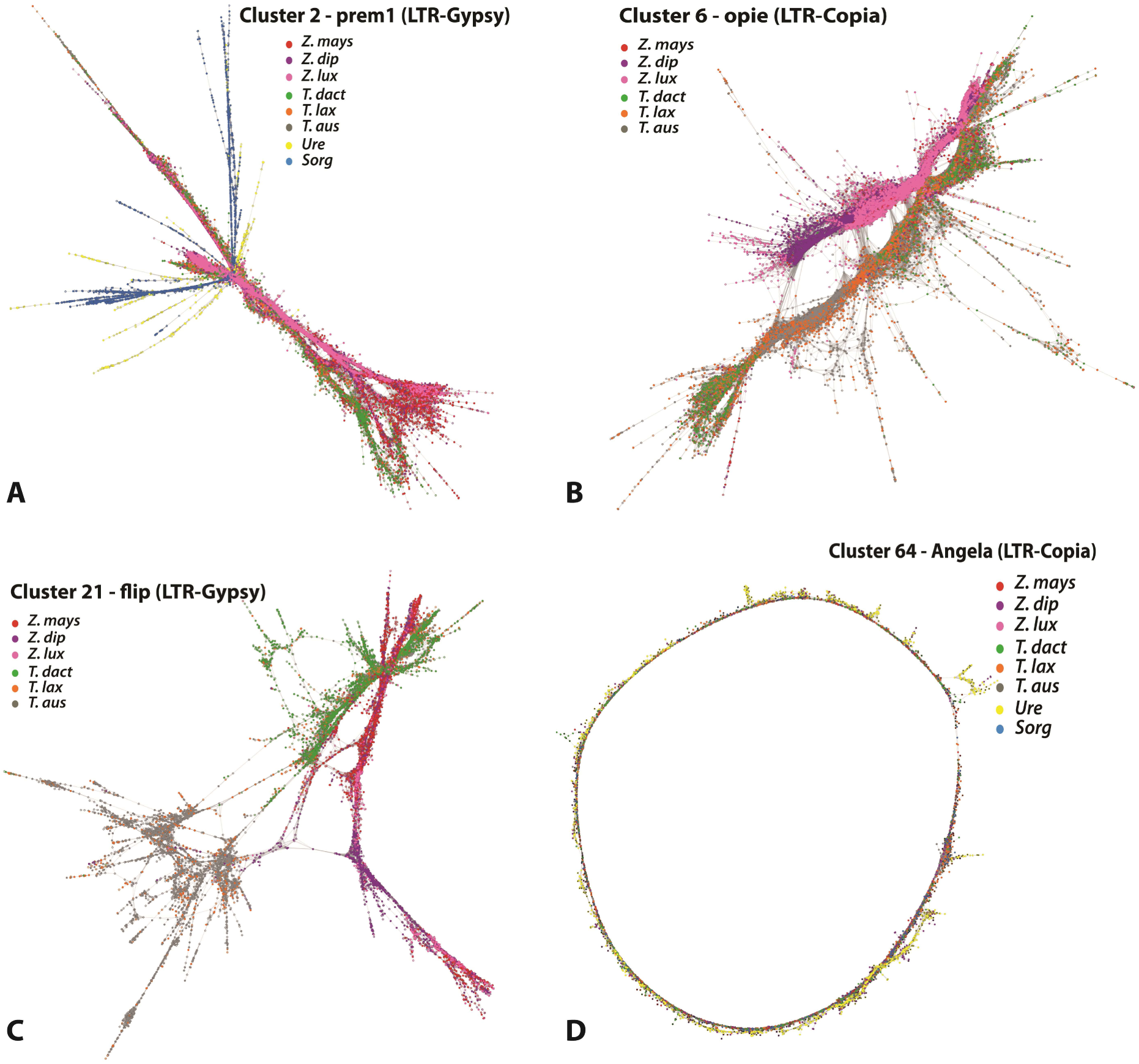
Comparative graph-based clustering. Graphs of individual repeat clusters that are shared between species demonstrating existing sequence variants within species. Highlighted dots represent sequences from individual species and lines connecting the dots represent sequence similarity. Each species is represented with a unique color: Red (*Z. mays*), purple (*Z. diploperennis*), pink *(Z. luxurians*), green (*T. dactyloides*), orange (*T. laxum*), and grey (*T. australe*). **A.** Cluster 2 shows shared LTR-*gypsy* elements in all genomes, in which sequences of *Zea* and *Tripsacum* are tightly connected with each other and sequences from *Urelytrum* and S*orghum* are peripherally connected, concordant with their evolutionary relationships. **B.** Cluster 6 (*Opie*, LTR-*copia*) is an example of a lineage-specific repeat family, where sequences are shared between *Zea* and *Tripsacum*; however, there is a clear separation in clustering of both lineages. **C.** Cluster 21 (*Flip*, LTR-*gypsy*) shows three separate groups in which the cultivated genomes *Z. mays* and *T. dactyloides* are more similar to one another than either is to their sister species. **D.** Cluster 64 (*Angela*, LTR-*copia*) is an example of a tightly knitted graph in a linear arrangement shared between all eight species, demonstrating the conserved nature of ancient *Angela* elements across all included taxa.

From a total of two million reads from eight genomes, 248 significant clusters were formed of various sizes and repeat families (**Figures 3A and B**). On average, ∼81% of the reads from each species clustered with LTR-retrotransposons (127 LTR-*gypsy* and 48 LTR-*copia* clusters). Among the 175 LTR-RT clusters (or families) identified, 85 families were present exclusively in the *Zea-Tripsacum* clade. For all species except *Z. mays* and *T. dactyloides*, the proportion of reads from LTR-*gypsy* families (53%) was higher compared to LTR-*copia* families (28%), whereas *gypsy* and *copia* were nearly equally abundant in *Z. mays* and *T. dactyloides* (**Figure 1A**). Compared to the other genomes, *Sorghum* contained the smallest proportion (11%) of reads from *copia* families.

**Figure 3.**
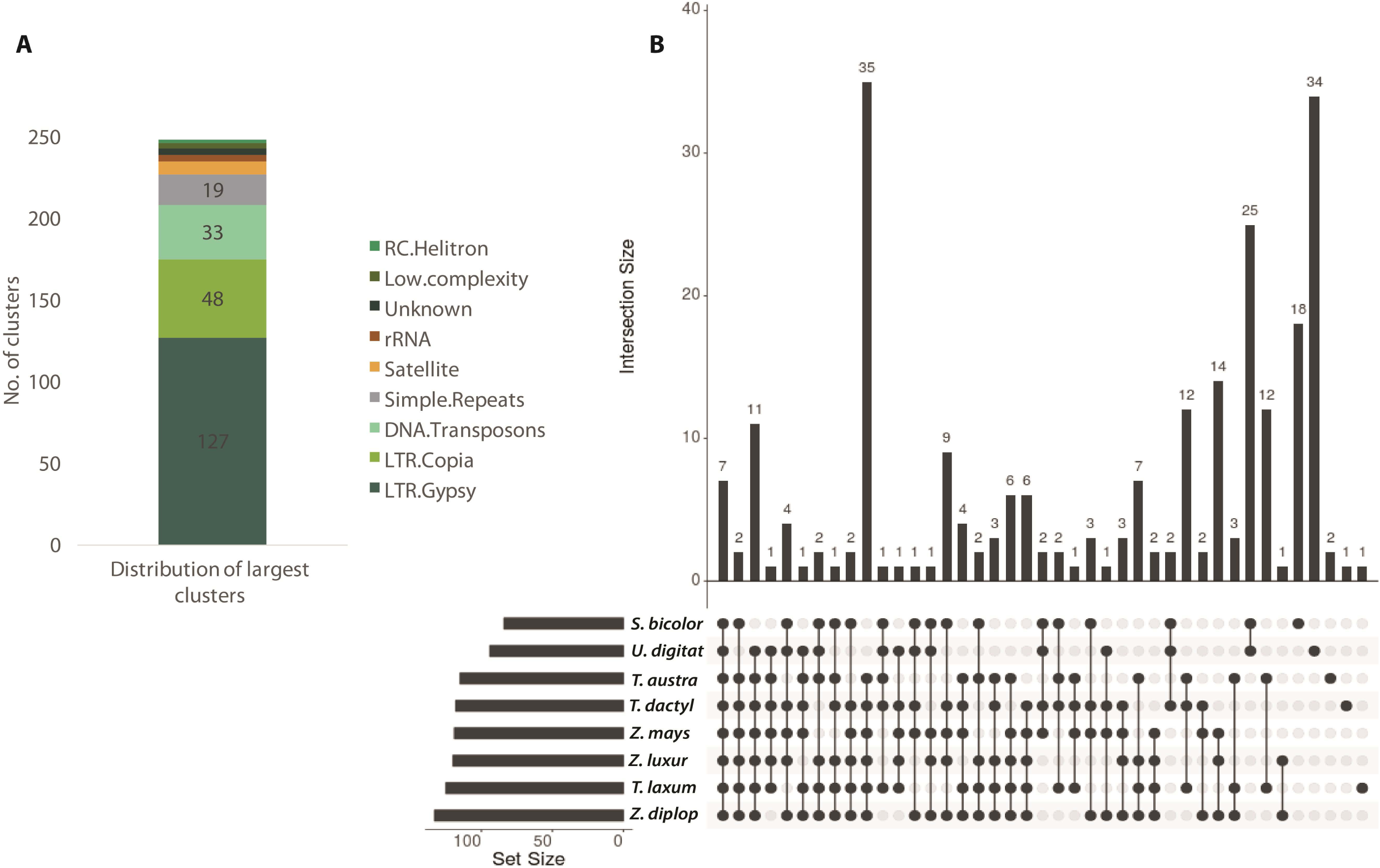
**A**. Bar graph showing the distribution of 248 largest clusters with respect to various repeat families. **B**. UpSet plot showing the interactions of shared repeat clusters among eight species. Each species is represented in one row with filled and empty cells. Each column represents the intersection between each species. From left to right, elements shared in all eight species to elements unique to each species is shown. Filled cells indicate that the element is shared with other species. The bars above each intersecting row represent the intersection size.

Among the 127 *gypsy* clusters, four clusters were shared among all eight species, two clusters were common to *Zea, Tripsacum*, and *Urelytrum* (but absent in *Sorghum*), 34 clusters were exclusive to *Zea* and *Tripsacum* species, and 15 clusters were found only in *Urelytrum* and *Sorghum* (**Figure S1A**). In addition, we observed lineage-specific *gypsy* families: ten in *Zea*, 17 in *Tripsacum*, 21 in *Urelytrum*, and 14 in *Sorghum*. Of the 48 *copia* clusters, only two were common to all species, ten were common to *Zea, Tripsacum* and *Urelytrum*, 19 clusters were exclusive to *Zea* and *Tripsacum*, and three were exclusive to *Urelytrum* and *Sorghum* (**Figure S1B**). Compared to *gypsy* super-families, there were fewer species-specific *copia* families.

### Insertional biases of *copia* elements in *Z. mays* and *T. dactyloides*

The *copia* superfamily was found to be more abundant in *Z. mays* and *T. dactyloides* compared to their relatives, suggesting recent lineage-specific proliferation of *copia* elements independently in both species. Investigating the genomic distribution of this expansion in *Z. mays*, we discovered that *copia* insertions were more frequently associated with stress-associated genes (34%) compared to those of other GO categories (on average 6%). *Copia* insertions mapped close to genes involved in plant defense, leaf morphogenesis, photoreceptors, homeobox proteins, signal transduction, and transcription (**Figure S2**). Further analysis revealed that these insertions are located in areas of high gene density, surrounded by approximately four to five genes within 5 kb windows both upstream and downstream. On repeating this analysis in *T. dactyloides* using the *Z. mays* reference protein coding genes, we discovered that the majority of *T. dactyloides copia* insertions are associated with genes involved in ubiquitin conjugation pathways, DNA metabolic processes, and macromolecule biosynthesis, in addition to stress and plant defense. In comparison, we did not find any close association between *S. bicolor copia* elements and stress-associated genes. Paired-end analysis revealed that the majority (∼85%) of *S. bicolor copia* reads were mapped to retrotransposon proteins and putative photoreceptor proteins. Unlike *Z. mays*, these elements were located in areas of lower gene density and surrounded by putative and unclassified genes.

### Evolutionary relationships and timing of transposition events

To assess the timing of major transposition events, we constructed maximum likelihood trees using the INT and RT domains (data not shown) of both *gypsy* and *copia* elements. Of the 127 shared *gypsy* clusters, 15 (total of 82 sequences) shared sufficient sequence identity within the integrase domain to allow amino acid sequence alignment. Major repeat families such as *Cinful-xeon, Prem1, Flip, Gyma, Xilon-Diguus*, and *Huck* were among these 15 clusters. With a few exceptions, most clades formed as expected in regard to species relationships. The *gypsy* families *Flip* and *Gyma* clustered together. The *Sorghum* and *Urelytrum* sequences from the *Flip* family clustered with *Zea* sequences of *Gyma*, whereas the *Zea* and *Tripsacum* sequences of *Flip* clustered with *Tripsacum* and *Sorghum* of *Gyma*. Several families such as *huck, puck*, and *grande* were clustered together with high support, suggesting a recent origin of these families. Clusters such as CL24 (unclassified), *uwum* (CL82) and *guhis* (CL132) also clustered with high sequence similarity.

We employed comparative sequence analyses of LTRs from 15 prominent clusters to estimate the temporal activity of retroelements both pre- and post-divergence of the *Zea-Tripsacum* clade (**Figure 4**). The clusters chosen for this analysis are composed of the following repeat families: *Prem1, Flip, Cinful-Xeon, Gyma, Ji, Opie, Dijap, Retrosor-6*, and several prominent unclassified elements. In **Figure 4A**, the peak activity of each element per species per cluster is plotted against a TE-specific grass molecular clock (11 mya to present). The approximate timing of the *Zea-Tripsacum* divergence is highlighted in yellow (5-6 mya). *Zea* and *Tripsacum* have experienced post divergence lineage-specific activity for most repeat families. For example, *Ji, Opie*, and *Dijap* (CL7, CL12, CL15, CL42, CL19, and CL51) were active within the last three million years for all species in which they are present. The *Opie* element represented in CL7 is shared between *Zea, Tripsacum*, and *Urelytrum* and has been active within the last ∼1-3 my indicating that amplification of *Opie* occurred in all three lineages (**Figure 4A and 4B**). In contrast, the amplification of CL2 (*prem1*) occurred recently only in *Z. luxurians* (2-3 mya) compared to all other species. Although *T. dactyloides* and *T. laxum* experience increased activity of *Prem1* around the time of divergence, the activity of this element in *Z. mays, Z. diploperennis, T. australe*, and *Sorghum* dates as an older amplification event. Similarly, the activity of CL5 (*gyma*) is recent in *Z. diploperennis* but older in *Z. luxurians*. Several families were shared only between *Sorghum* and the wild relatives of *Zea (Z. diploperennis, Z. luxurians)* and *Tripsacum (T. laxum, T. australe)*. Despite their presence, the activity of these families varies among species. For example, the activity of CL11 in *Z. luxurians* is recent (0-1 mya) but is dated as an older event in the other species (**Figure 4A and 4C**).

**Figure 4.**
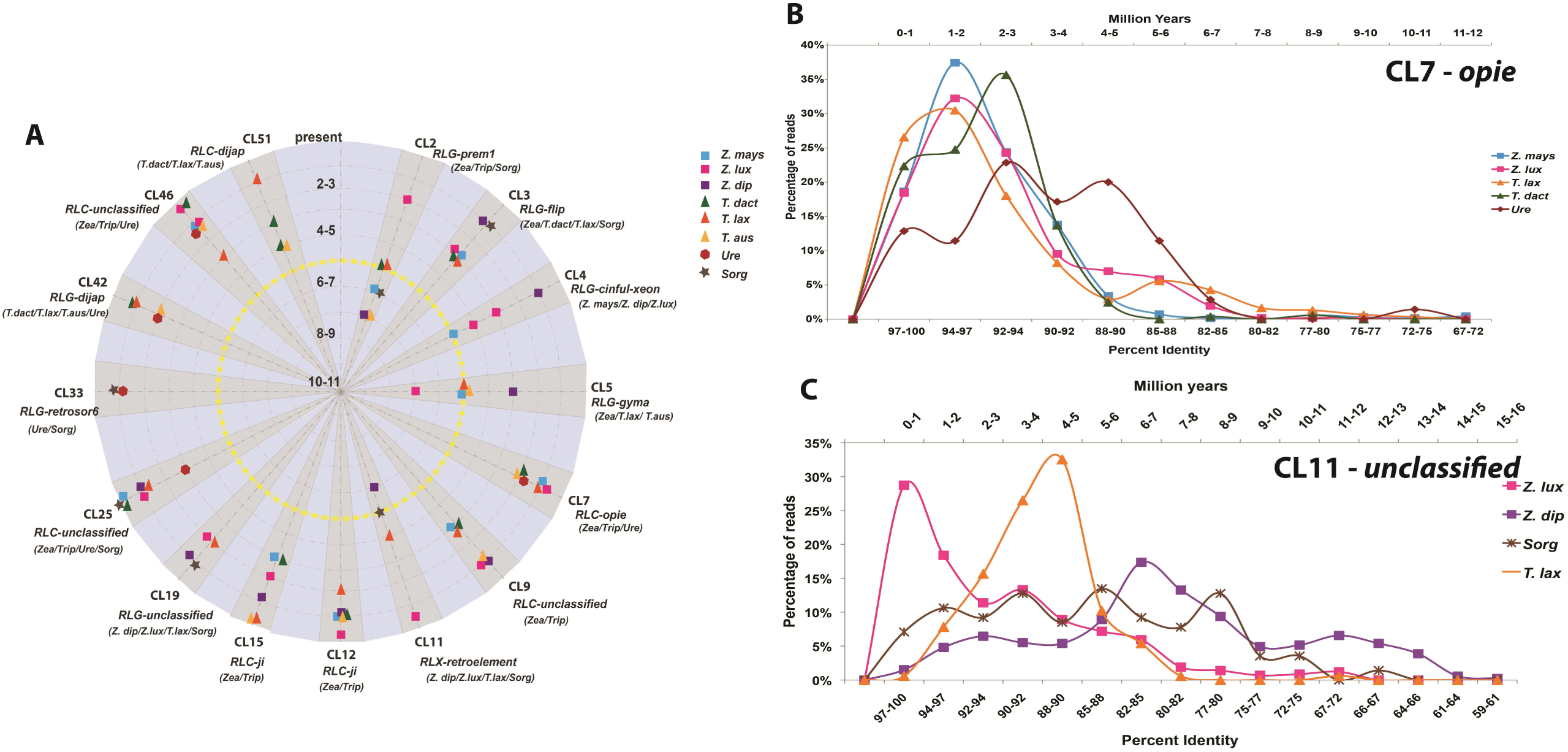
**A**. Activity of retroelements pre-and post-divergences of the *Zea-Tripsacum* clade (yellow line) for 15 prominent retrotransposon clusters (shaded gray). Concentric circles indicate time scale per million years from 11 mya (center) to present (outer circle). For each cluster, the corresponding repeat family and shared species information is given below each cluster name. Each data point represents the peak activity of that element. **B & C** display retrotransposon activity of CL7 & CL11 based on percent identity of shared LTR sequences (bottom axis) and the corresponding grass molecular clock (mya) along the top axis. CL11 is absent in domesticated *Zea* and *Tripsacum* but present in the wild relatives.

## Discussion

The present study evaluates TE dynamics in divergent descendants (*Zea* and *Tripsacum*) of a common allopolyploid ancestor within a phylogenetic framework and in comparison to two diploid relatives (*Urelytrum digitatum* and *Sorghum bicolor*). The comparative analysis of repeat elements from *Zea, Tripsacum, Urelytrum*, and *Sorghum* provides insight into the contribution of retrotransposons to genome evolution after a shared polyploidization event. Inclusion of additional *Zea* and *Tripsacum* species provided an opportunity to assess the genomic variability in repeat content among species within a genus.

As expected, LTR-retrotransposons account for the majority of the repeats in the genomes of all species included in this study. Individual clustering analyses indicate that diverse LTR-retrotransposons contribute to genome size variation in this taxonomic group. Based on our comparative and molecular clock analyses, the majority of retrotransposon families are common to the *Zea – Tripsacum* clade in comparison to their diploid relatives, suggesting the occurrence of retrotransposon proliferation after allopolyploidization but before the split between the two genera (**Figure 4A**). Previous studies have hypothesized an occurrence of retroelement bursts just before the divergence of *Zea* and *Tripsacum* based on maize retroelement activity (Gaut et al. 2000, Estep et al. 2013). The results for the *Tripsacum* species included in the current analysis support this hypothesis, revealing a high number of shared retrotransposon families between the two genera. For example, *Ji* and *Opie* of the *copia* superfamily have been especially active (2 mya, **Figure 4A**, >300 Mb, **Table 1**) in both *Zea* and *Tripsacum*; however, these families contribute little (∼35 Mb) to genome composition in *Urelytrum* and are absent in *Sorghum*. The presence and hyperactivity of these families in *Zea* and *Tripsacum* but not in *Sorghum* suggests amplification after the maize-sorghum split but before the allopolyploidization event leading to the *Zea-Tripsacum* lineage. Similarly, five *gypsy* families (*Cinful-Xeon, Prem1, Flip, Gyma*, and *Huck*) are abundant in the *Zea-Tripsacum* clade but present in low copy numbers in the other lineages. Molecular clock analysis reveals recent activity (1-4 mya) for these families in *Zea* and *Tripsacum*, suggesting amplification after divergence from other taxa (**Figure 4A**). Conversely, families such as *Athila* and *Leviathan* have accumulated in the *Urelytrum* and *Sorghum* genomes, but are absent in *Zea* and *Tripsacum*, suggesting independent activation of LTR-retrotransposon families in different lineages over short evolutionary time scales.

Additionally, both *Zea* and *Tripsacum* contain lineage-specific families, indicating variation in retroelement amplification in each genus after divergence. Compared to *Zea, Tripsacum* contained more unique *gypsy* and *copia* families. Nine *gypsy* families were common to all *Tripsacum* genomes and seven families were shared only between *T. laxum* and *T. australe* (**Figure 3B & Figure 4A)**. The larger number of unique and recently active retroelements (∼1-4 mya) in *Tripsacum* indicates the independent expansion of these families after the *Zea-Tripsacum* divergence. Overall, the abundance and recent activity of these genus-specific LTR-retrotransposons shared only between *Zea* and *Tripsacum* suggests that the activation of these families might be an outcome of shared polyploidization as proposed by the genomic shock hypotheses (McClintock 1984, Comai et al. 2003).

Surprisingly, clustering analyses suggest that *Z. mays* and *T. dactyloides* share greater similarity in TE composition and abundance for some retrotransposon families than to the other members of their respective genera. Examples of noticeably increased abundance in these two species include the families *Ji, Flip*, and *Opie*. Conversely, we also found significantly decreased copy numbers of *Huck, Doke*, and *Puck*. Also, a greater number of reads from both genomes are derived from the *copia* superfamily, indicating independent expansion of *copia* clades in both lineages, similar to results from comparisons in Asian rice varieties (Li et al. 2017). For a few shared *copia* clusters, the peak of activity in *Z. mays* overlaps with *T. dactyloides*, suggesting both species experienced *copia* activity during a similar time period (**Figure 4A)**. Considering the independent evolution of both species post divergence and the role of artificial selection in maize domestication, the similarity in composition and activity of *copia* elements in *Z. mays* and *T. dactyloides* suggests that natural and artificial selection have influenced TE amplification and accumulation similarly in both lineages. Although the precise cause of such species-specific activity is unclear, we propose that the observed patterns in TE abundance and accumulation may be related to adaptation to temperate climates. Indeed, recent evidence shows parallels in protein sequence evolution between natural and artificial selection in *T. dactyloides* and maize during their adaptation to temperate climates (Yan et al. 2019).

Additionally, considering the intensity of retroelement accumulation in maize as reported in other studies, it is likely that these *copia* elements have been active during maize domestication and improvement. Studies demonstrating TE involvement in plant domestication predominantly show that insertions near functional genes play a role in plant function and/or development. Well-known examples include *Hopscotch* involvement in apical dominance in maize (Studer et al, 2011), *Gret1* in berry color variation in *Vitis vinifera* (Cadle-Davidson et al. 2008), and LTR-mediated control of the blood orange phenotype (Butelli et al. 2012). Because the clustering analysis revealed recent *copia* expansion in both *Z. mays* and *T. dactyloides*, we explored the frequency of *copia* insertions near genes. In *Z. mays*, a large number of reads from *copia* elements mapped close to several functionally relevant genes. Indeed, the majority of *copia*-associated genes are involved in plant defense, homeobox proteins responsible for shoot apical meristem and leaf morphogenesis, cytokinin response, signal transduction, and transcription (**Figure S2**).

Similar functional gene categories were associated with *T. dactyloides copia* elements; however the majority of the insertions were associated with genes involved in stress associated pathways such as ubiquitin conjugating pathway and macromolecule biosynthesis pathways, including carbohydrates, lipids, and proteins. Additionally, although gene density in *Z. mays* is approximately one gene per 3.2 kb (Fu et al. 2001), the *copia* insertions identified in this study were surrounded by approximately 4-5 genes upstream and downstream in a 5kb interval, suggesting biased insertion in more gene-rich regions. Such close proximity provides the potential for TEs to affect the function of neighboring genes, as seen in other plants (Makarevitch et al. 2015, Cao et al. 2016, Pietzenuk et al. 2016). For example, the tobacco *Tnt1* and the rice *Tos17 copia* elements were found near stress-related genes, and the expression of these elements is linked with the biological responses of the plant to the external stresses (Grandbastein et al. 1997, Miyao et al. 2003, Le et al. 2007).

To test whether this pattern is common to domesticated plants, we performed the same analysis for *S. bicolor*. Although both *Z. mays* and *S. bicolor* are closely related grasses, we did not find similar *copia* insertions near genes that are involved in *S. bicolor* plant development and defense. For example, in *Z. mays*, 32 *copia* elements were inserted within 1kb of a stress gene (GRMZM2G047919) whereas in *S. bicolor* only two *copia* elements were found near a stress related gene (SORBI_3009G188300). We suggest that the observed *copia* expansion is therefore unrelated to the process of domestication, but rather a byproduct of genome duplication. In other words, the disruptive nature of insertional mutagenesis may be buffered in *Z. mays* and *T. dactyloides* by the presence of duplicate genes in the allopolyploid.

## Conclusion

In this study, we provide insight into interspecific TE diversity and its contribution to genome evolution in related members of Andropogoneae that have undergone a shared polyploidization event. By including multiple accessions of two divergent species (*Zea* and *Tripsacum*) originating from a common allopolyploid ancestor, in addition to close diploid relatives (*Urelytrum digitatum* and *Sorghum bicolor*), we described LTR-retrotransposon diversity with respect to the hybridization and genome doubling process. Though the genome size of *Urelytrum* is similar to that of *Sorghum*, the repeat composition of *Urelytrum* is more like that of *Zea* and *Tripsacum*. Similarities in the proportion of the *copia* superfamily and satellite DNA in *Zea-Tripsacum-Urelytrum* suggests that *Urelytrum* or a close relative played an ancestral role in the origin of the *Zea-Tripsacum* lineages, supporting the hypothesis proposed by McKain et al. (2018). Our clustering analysis revealed an expansion of the *copia* superfamily exclusively in *Z. mays* and *T. dactyloides*, suggesting amplification (and possibly participation) of new *copia* insertions during adaptation to temperate environments. Indeed, many of the insertions are near genes involved in plant development, defense, and macromolecule biosynthesis. Therefore, the *cis*-regulatory effects of TE insertions near genes may have influenced the evolution of *Z. mays* and *T. dactyloides* during both adaptation and domestication.

## Acknowledgements

We acknowledge the use of the Super Computing System Spruce Knob at West Virginia University, which is funded by the National Science Foundation EPSCoR Research Infrastructure Improvement Cooperative Agreement 1003907, the state of West Virginia (WVEPSCoR via the Higher Education Policy Commission) and West Virginia University.

## Figure legends

**Figure S1.**
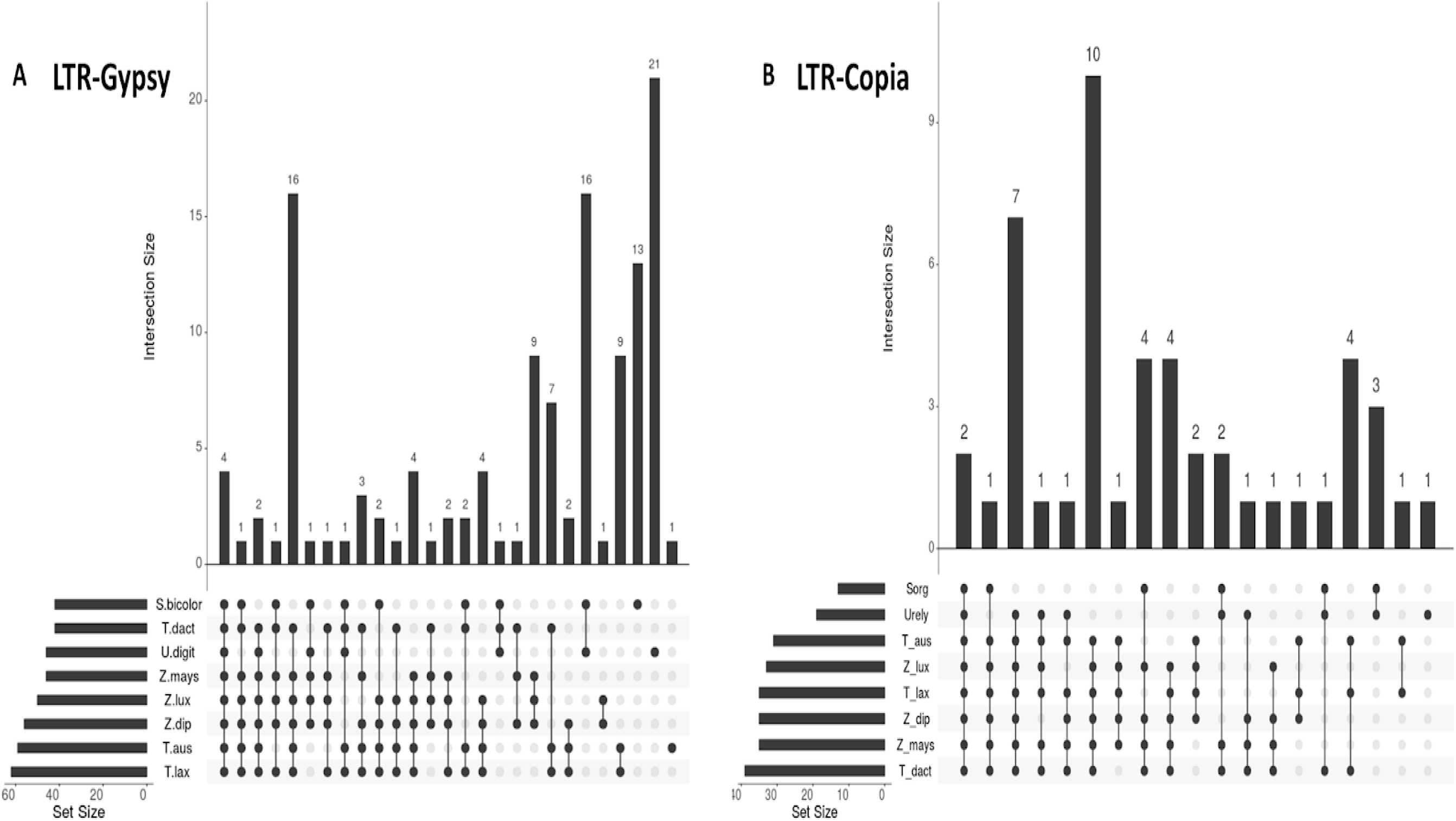
UpSet plots showing the interactions of shared *gypsy* (A) and *copia* (B) clusters. Each species is represented in one row with filled and empty cells. Each column represents the intersection between each species. Clusters are displayed from most common (on the left) to least common (on the right). Cells are either filled or empty indicating whether the element is shared with other species. The bars above each intersecting row represent the intersection size.

**Figure S2:**
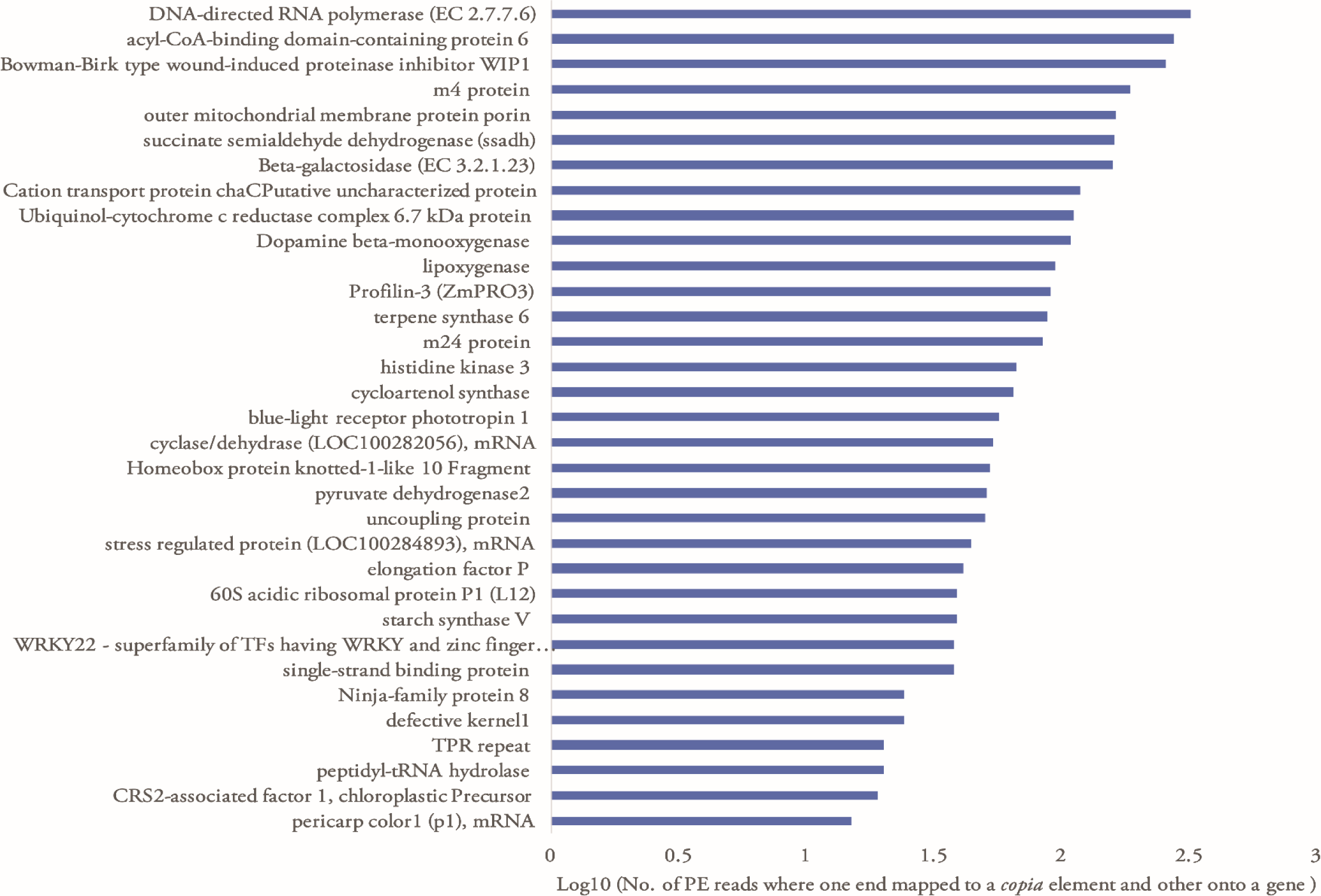
Paired-end read mapping to *copia* elements and nearby gene in *Z. mays.* The majority of the TEs proximal to genes are involved in plant development and defense, such as terpene synthase, beta galactosidase, profilin-3, and blue-light receptor phototropin.

